# Receptive field center-surround interactions mediate context-dependent spatial contrast encoding in the retina

**DOI:** 10.1101/252148

**Authors:** Maxwell H. Turner, Gregory W. Schwartz, Fred Rieke

## Abstract

Antagonistic receptive field surrounds are a near-universal property of early sensory processing. A key assumption in many models for retinal ganglion cell encoding is that receptive field surrounds are added only to the fully formed center signal. But anatomical and functional observations indicate that surrounds are added before the summation of signals across receptive field subunits that creates the center. Here, we show that this receptive field architecture has an important consequence for spatial contrast encoding: the surround can control sensitivity to fine spatial structure by changing the way the center integrates visual information over space. The impact of the surround is particularly prominent when center and surround signals are correlated, as they are in natural stimuli. This role of the surround differs substantially from classic center-surround models and raises the possibility that the surround plays unappreciated roles in shaping ganglion cell sensitivity to natural inputs.

## Introduction

The receptive field (RF) surround is a ubiquitous feature of early sensory computation. Surrounds in the retina have been proposed to enhance sensitivity to edges in a visual scene via lateral inhibition [1]. Surrounds have also been suggested to promote efficient representation of naturalistic visual stimuli. Natural scenes contain strong correlations across visual space [2, 3, 4], and the RF surround can decorrelate responses across a population of visual neurons [5, 6, 7], thereby increasing encoding efficiency by reducing redundancy in the population code [8, 9].

Recent findings have challenged this proposed function of the RF surround by showing that mechanisms other than the surround can play a larger role in decorrelating (or “whitening”) natural images. First, nonlinear processing in the retina can contribute more to decorrelating retinal ganglion cell (RGC) responses to natural scenes than the RF surround [10] (see also [11, 12]). Second, human fixational eye movements can remove spatial correlations in natural inputs before any neural processing takes place [13, 14, 15, 16]. These findings suggest a need to re-examine the role of the surround, especially as it relates to the encoding of natural visual stimuli.

Here, we answer several questions about center-surround RF structure and its relation to natural scenes. First, how does the circuit location of surround suppression (before or after summation over visual space) impact RGC computation? Second, does surround activation change the linearity of spatial integration by RGCs? And finally, how do the statistics of natural scenes influence nonlinear interactions between the center and the surround?

Classical Difference-of-Gaussians [17, 18, 19, 20] and modern predictive RF models [21, 22] assume that the surround interacts only with the fully formed center signal (Fig. 1A). These models suggest that the surround modulates the gain of signals in the center but does not alter how the center integrates signals across space. However, surrounds are generated by horizontal cells in the outer retina [23, 24, 25, 26, 27] and amacrine cells in the inner retina [28, 29, 30, 31, 32] (Fig. 1C). As a result, at least a portion of the surround is present in the bipolar cells presynaptic to a RGC (Fig. 1C, S1;[33, 34, 35, 36, 11, 37]) and hence is present prior to the summation over space that forms the center. Subunits with antagonistic surrounds have been incorporated into models of the RGC RF [38]. Nonetheless, the RF structure in Fig. 1B is not reflected in modern, predictive models nor have its functional consequences been explored.

**Figure 1:**
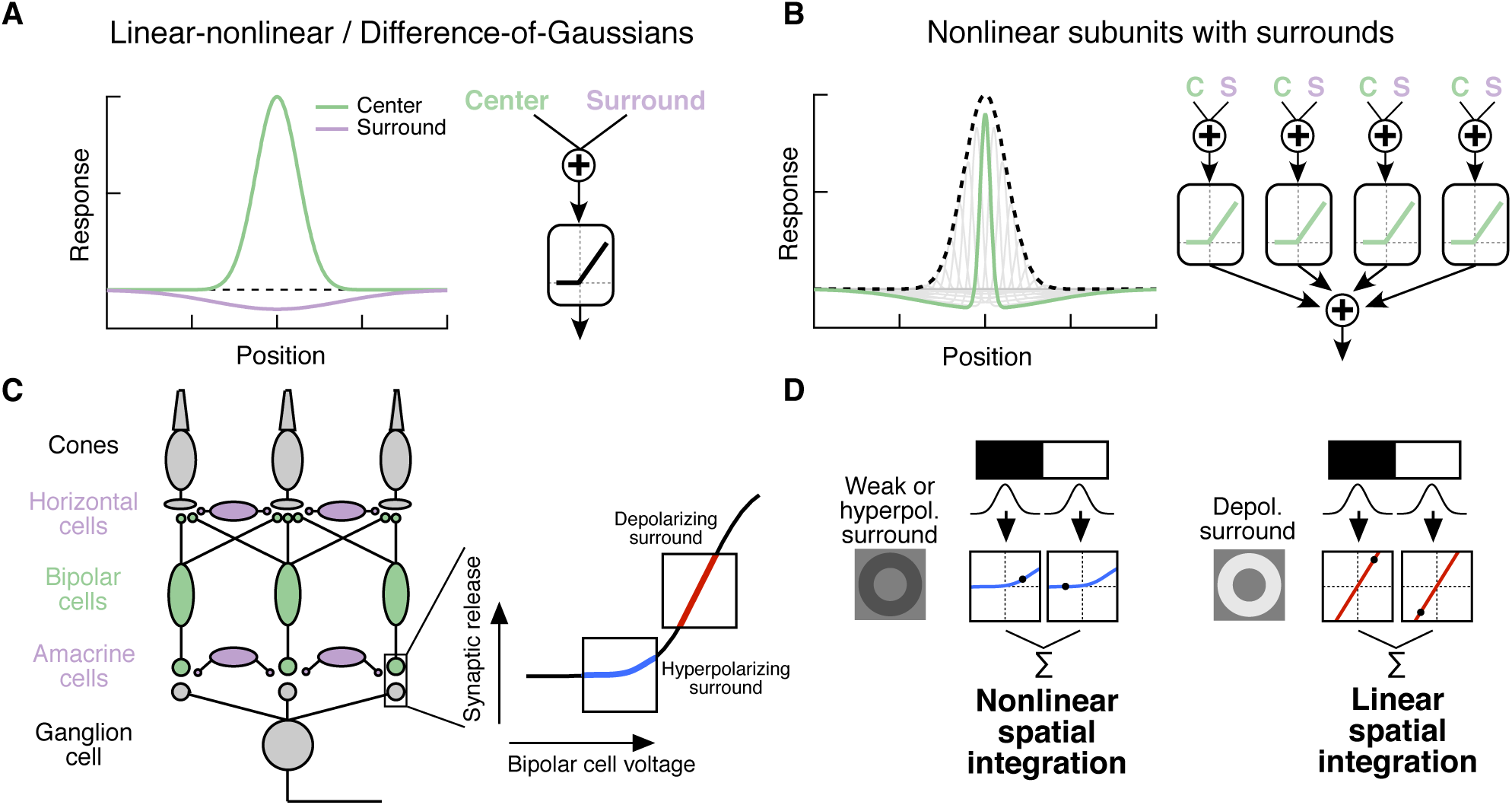
Circuit basis of the surround and implications for receptive field structure. (A,B) Two competing models for how the surround is integrated with the center to form the full RF: The linear-nonlinear model shown in (A) linearly combines the center and surround before treatment with an output nonlinearity that occurs after spatial integration. The model in (B) combines center and surround signals within each nonlinear subunit, and treats the combined center-surround signal with a private nonlinearity. In other words, individual subunits in (B) have their own RF surrounds. (C) Retinal circuit schematic. Bipolar cells make excitatory synapses on to retinal ganglion cells (RGCs). The nonlinear synaptic transfer from bipolar cell to RGC means that bipolar cells act as nonlinear subunits within the receptive field (RF) center of the RGC. The surround is generated by horizontal and/or amacrine cells, both of which can influence bipolar cell responses upstream of the synaptic nonlinearity. A bipolar-cell hyperpolarizing surround input shifts the synapse into a more rectified state (blue portion of synaptic nonlinearity). A depolarizing surround input shifts the synapse into a locally-linear state (red portion of curve). (D) Based on this circuit organization, we hypothesized a working model of how inputs to the surround could change RF structure in the center. Shown is a schematic illustrating this hypothesis for an Off-center RGC. A weak or hyperpolarizing surround will be associated with rectified subunit output nonlinearities, and thus nonlinear spatial integration and sensitivity to spatial contrast (e.g. a split-field grating stimulus). A depolarizing surround will be associated with more linear subunit outputs and the RGC should integrate across visual space approximately linearly.

For many RGC types bipolar cells act as excitatory, nonlinear subunits within the RF center [39, 40] (Fig. 1C). The degree of rectification at the bipolar cell to RGC synapse can control the spatial integration properties of the RF center [41, 42]. In particular, a weakly rectified or linear synapse yields linear spatial integration due to cancellation of bright and dark regions of space, while a sharply rectified bipolar output synapse yields nonlinear spatial integration because light and dark regions fail to cancel. Nonlinear spatial integration causes RGCs to respond strongly to stimuli with fine spatial structure like gratings [43, 44, 18, 45, 46].

We hypothesized that surround modulation could dynamically regulate nonlinear spatial integration in a RGC by controlling the effective rectification of the bipolar synapse. Thus, in response to weak surround input or surround input that hyperpolarizes bipolar cells, bipolar cell terminals presynaptic to the RGC are in a rectified state and the RF center integrates nonlinearly over visual space (Fig. 1C,D). When surround input depolarizes bipolar cells in the RF center, however, the bipolar cell synapse is relatively more linear, and the RF center integrates linearly over visual space (Fig. 1C,D).

We test this hypothesis using both synthetic and natural visual stimuli to probe responses of parasol (magnocellular-projecting) RGCs in the macaque monkey retina. The nonlinear subunit structure of these cells has been well characterized [43, 44, 46], and this RF organization can be important for encoding natural visual stimuli [42]. Our results show that RGC sensitivity to spatial contrast in natural scenes is modulated by context via a nonlinear interaction between the RF center and surround.

## Results

We use single cell patch electrophysiology in an *in vitro* macaque retinal preparation in conjunction with computational modeling of the RGC RF to explore the impact of the surround on nonlinear RF structure and natural scene encoding. We start by testing the hypothesis outlined in Fig. 1C,D. We find that, indeed, spatial integration in the RF center depends strongly on surround activation. Next, we use natural and artificial stimuli to characterize nonlinear interactions between center and surround and test circuit models for the origin of these interactions. Finally, we show that the intensity correlations characteristic of natural scenes promote nonlinear interactions and make spatial integration relatively insensitive to changes in local luminance across a visual scene.

### The RF surround regulates nonlinear spatial integration in the RF center

We focused our test of the hypothesis in Fig. 1C,D on Off parasol RGCs, because these cells show both stronger rectification of subunit output than On parasol cells [47] and nonlinear spatial integration in the context of naturalistic visual stimuli [42]. We began each experiment by centering the stimulus over the RF and measuring the linear RF (see methods and Fig. S1). We then tailored visual stimuli for each cell such that the “center” stimulus did not extend into the pure surround RF subregion and the “surround” stimulus did not cover the center. This allowed us to ask questions about interactions between these two RF subregions. These subregions are not exclusively associated with a center or surround mechanism since the antagonistic surround is spatially coextensive with the RF center (Fig. 1A,B). Nonetheless, according to estimated RFs, the “center” stimulus activated the center mechanism *∼*4 times more strongly than the surround mechanism and the “surround” stimulus activated the surround mechanism *∼*5 times more strongly than the center mechanism.

We previously found that nonlinear spatial integration endows Off parasol RGCs with sensitivity to spatial contrast in natural images [42]. To test whether surround activity modulates this spatial contrast sensitivity, we measured Off parasol RGC spike responses to natural image patches that contain high spatial contrast and were expected to activate the nonlinear component of the cell’s response (see Methods for details on images and patch selection). For each image patch, we also presented a linear equivalent disc stimulus, which is a uniform disc with intensity equal to a weighted sum of the pixel intensities within the RF center. The weighting function was an estimate of each cell’s linear RF center from responses to expanding spots (see methods and Fig. S1 for example fits). A cell whose RF center behaves according to this linear RF model will respond equally to a natural image and its associated linear equivalent disc.

As shown previously, Off parasol cells responded much more strongly to natural images than to linear equivalent stimuli, especially when the natural image contains high spatial contrast (Fig. 2A, left, see also [42]). However, when a bright surround was presented with the center stimulus, the natural image and its linear equivalent disc produced near-equal responses (Fig. 2A,B). We used the difference in spike count between a natural image and its linear equivalent disc as a metric of spatial contrast sensitivity. This difference, as shown in Fig. 2C, depended systematically on the difference in mean intensity in the center and surround regions. Specifically, spatial contrast sensitivity was maximal in response to stimuli for which center and surround intensities were similar and dropped as intensity in these regions diverged. Hence, nonlinear spatial integration is maximized when center and surround experience similar mean luminance. When the surround strongly hyperpolarizes presynaptic bipolar cells (for Off cells, a dark surround), the response is diminished as a result of the surround shifting the synapse into a quiescent state. When the surround depolarizes presynaptic bipolar cells (for Off cells, a bright surround), the nonlinear sensitivity of the center is reduced, as outlined in Fig. 1.

**Figure 2:**
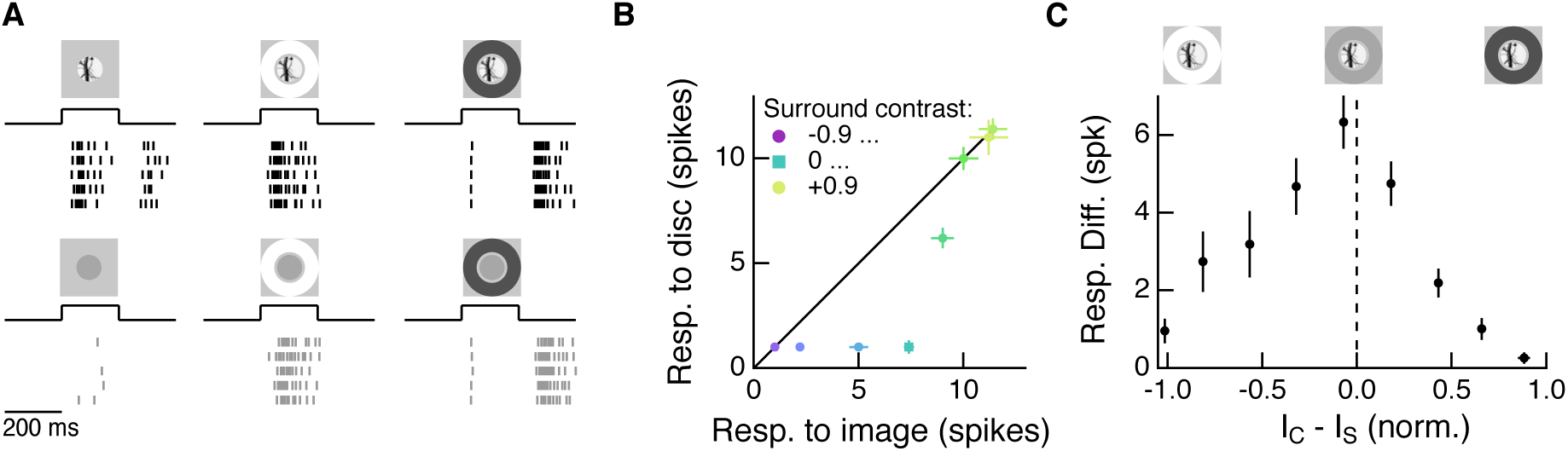
The RF surround regulates nonlinear spatial integration of natural images. (A) We presented a natural image patch and its linear-equivalent disc stimulus to probe sensitivity to spatial contrast in natural scenes. Example Off parasol RGC spike responses are shown. (B) Spike count responses to an example image patch and its linear equivalent disc across a range of surround contrasts. The addition of a sufficiently bright surround (top three points) eliminates sensitivity to spatial contrast in this image patch. (C) Population summary showing the response difference between image and disc as a function of the difference in mean intensity between the RF center and surround. Negative values of this difference correspond to a surround that is brighter than the center, and positive values to a relatively darker surround (n = 21 image patch responses measured in 5 Off parasol RGCs).

Does the impact of the surround on spatial integration in the center depend on specific statistics of natural images or is it a more general phenomena? To answer this question, we repeated these experiments using a flashed split-field grating rather than natural image patches (Fig. 3). Because of the nonlinear subunit structure of the RF center, Off parasol cells respond strongly to such stimuli (Fig. 3A). We presented the same grating together with a surround annulus and, in separate trials, the surround annulus alone (Fig. 3A). When paired with a bright surround, the grating stimulus did not activate the cell much beyond its response to the surround alone. A dark surround suppressed responses with and without the grating, save for a small, brief response at the beginning of the presentation of the grating which is likely the result of the brief temporal delay of the surround relative to the center.

**Figure 3:**
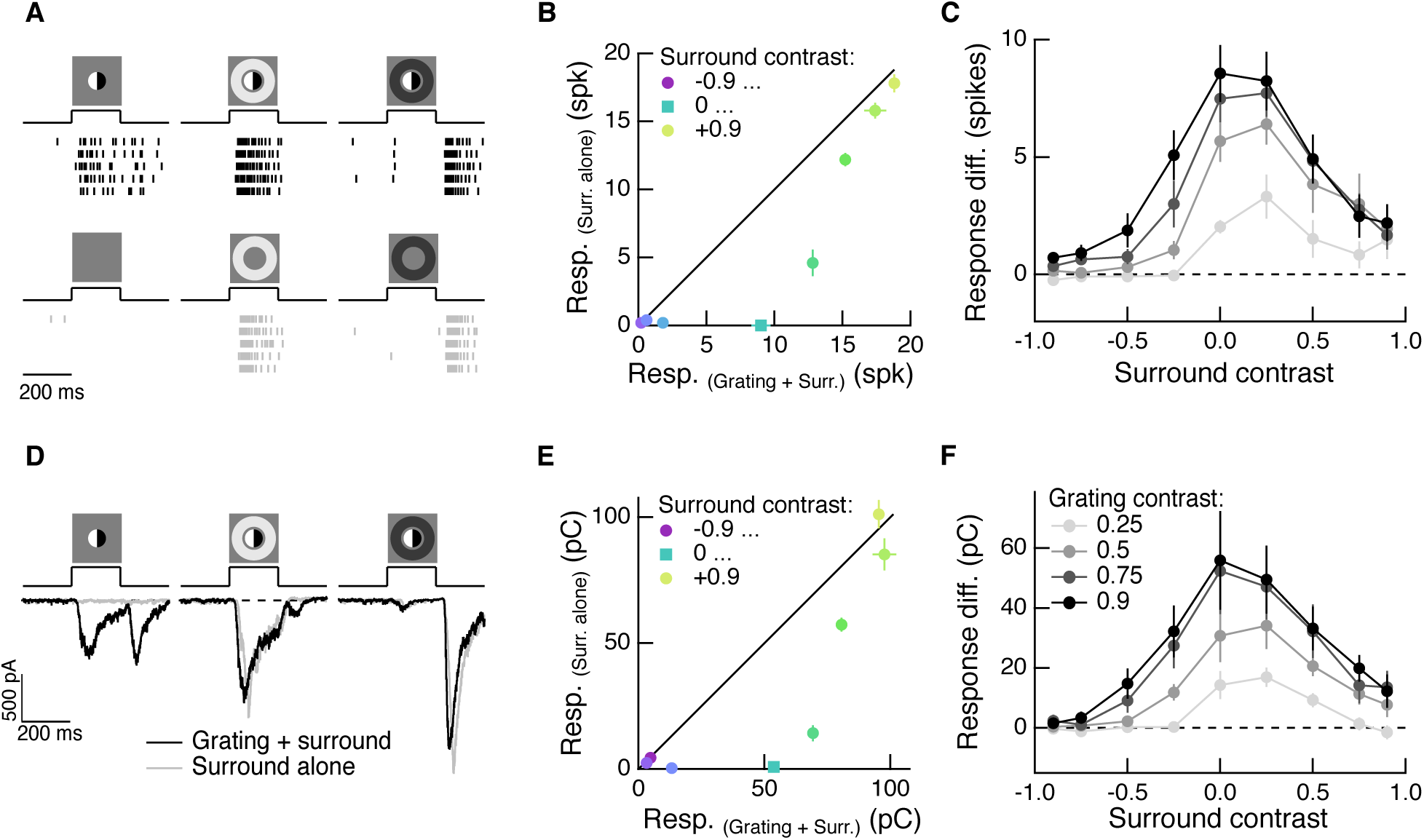
The RF surround regulates nonlinear spatial integration in the RF center. (A) Left column: Example Off parasol RGC spike response to an isolated split-field grating stimulus in the RF center. The linear equivalent stimulus (i.e. no stimulus) causes no response in the cell. Center column: when the center stimulus is paired with a bright surround, the grating and the linear equivalent stimulus produce very similar spike responses. Right column: a dark surround suppresses the response in both cases, and the grating is unable to elicit a strong response at all. (B) For the example cell in (A), we tested sensitivity to the center grating stimulus with a range of contrasts presented to the surround. Negative contrast surrounds (hyperpolarizing for off bipolar cells) decrease the response. Positive contrast surrounds (depolarizing for off bipolar cells) sum sub-linearly with the grating stimulus such that for the brightest surrounds, the addition of the grating only mildly enhances the cell’s response. Points show mean *±* S.E.M. spike count. (C) We measured the response difference between the grating stimulus and the surround-alone stimulus across a range of surround contrasts (horizontal axis) and for four different central grating contrasts (different lines). For each grating contrast, addition of either a bright or dark surround decreased sensitivity to the added grating. Points are population means *±* S.E.M. (n = 5 Off parasol RGCs). (D-F) same as (A-C) for excitatory synaptic current responses of an Off parasol RGC. Points represent mean *±* S.E.M. excitatory charge transfer for the example cell in (E) and population mean *±* S.E.M. (n = 7 Off parasol RGCs) in (F).

We repeated this experiment for surround contrasts ranging from +0.9 to −0.9 (Fig. 3B). While the surround-free grating (surround contrast = 0) stimulus showed strong nonlinear integration (indicated by its distance away from the unity line in Fig. 3B), the presence of a surround stimulus diminished the response to the grating (indicated by the tendency of non-zero surround contrast points to lie closer to the line of unity). This was true for a range of central grating contrasts (Fig. 3C), which suggests that this behavior is not the result of response saturation. Thus, just as for natural image patches, nonlinear spatial integration is maximal when the center and surround experience the same mean luminance (in this case a mean of zero) and decreases when the surround is brighter or dimmer than the center.

To test whether the effect of the surround on spatial integration was present in the bipolar synaptic output, we repeated these experiments while measuring a ganglion cell’s excitatory synaptic inputs (Fig. 3D; see methods for isolation of excitatory inputs). Modulation of spatial integration by surround activity was similar in excitatory inputs and spike responses (Fig. 3E,F), which suggests that it is not substantially shaped by inhibitory input, post-synaptic integration, or spike generation mechanisms. A similar effect can be seen in the excitatory inputs to On parasol RGCs (Fig. S2). Compared to Off parasol RGCs, On parasol RGCs are more easily shifted into a regime of linear spatial integration, presumably because of the shallower rectification of nonlinear subunits in the receptive field of On parasol cells [47, 42].

The experiments described in Figs. 2 and 3 show that inputs to the RF surround can influence how the RF center integrates signals across space, consistent with the hypothesis outlined in Figure 1. For both the spike output and excitatory synaptic inputs to Off parasol RGCs, the peak spatial nonlinearity was observed when center and surround experienced similar mean luminance (Fig. 2C and Fig 3C,F).

### Nonlinear center-surround interactions are dominated by a single, shared nonlinearity

The hypothesis in Figure 1 relies on a specific form of nonlinear center-surround interaction, whereby center and surround signals combine upstream of a shared, rectifying nonlinearity. To characterize center-surround interactions in an unbiased manner, we used Gaussian-distributed white noise stimulation and a linear-nonlinear cascade modeling approach. We presented Gaussian-distributed random noise to the center, surround or both regions while measuring excitatory synaptic inputs to On and Off parasol cells (Fig. 4A). For this analysis, we estimated the excitatory conductance by dividing the measured excitatory currents by the driving force. We computed linear filters for each RF region using reverse correlation based on trials in which the center or surround was stimulated in isolation (Fig. 4A, left and middle columns).

**Figure 4:**
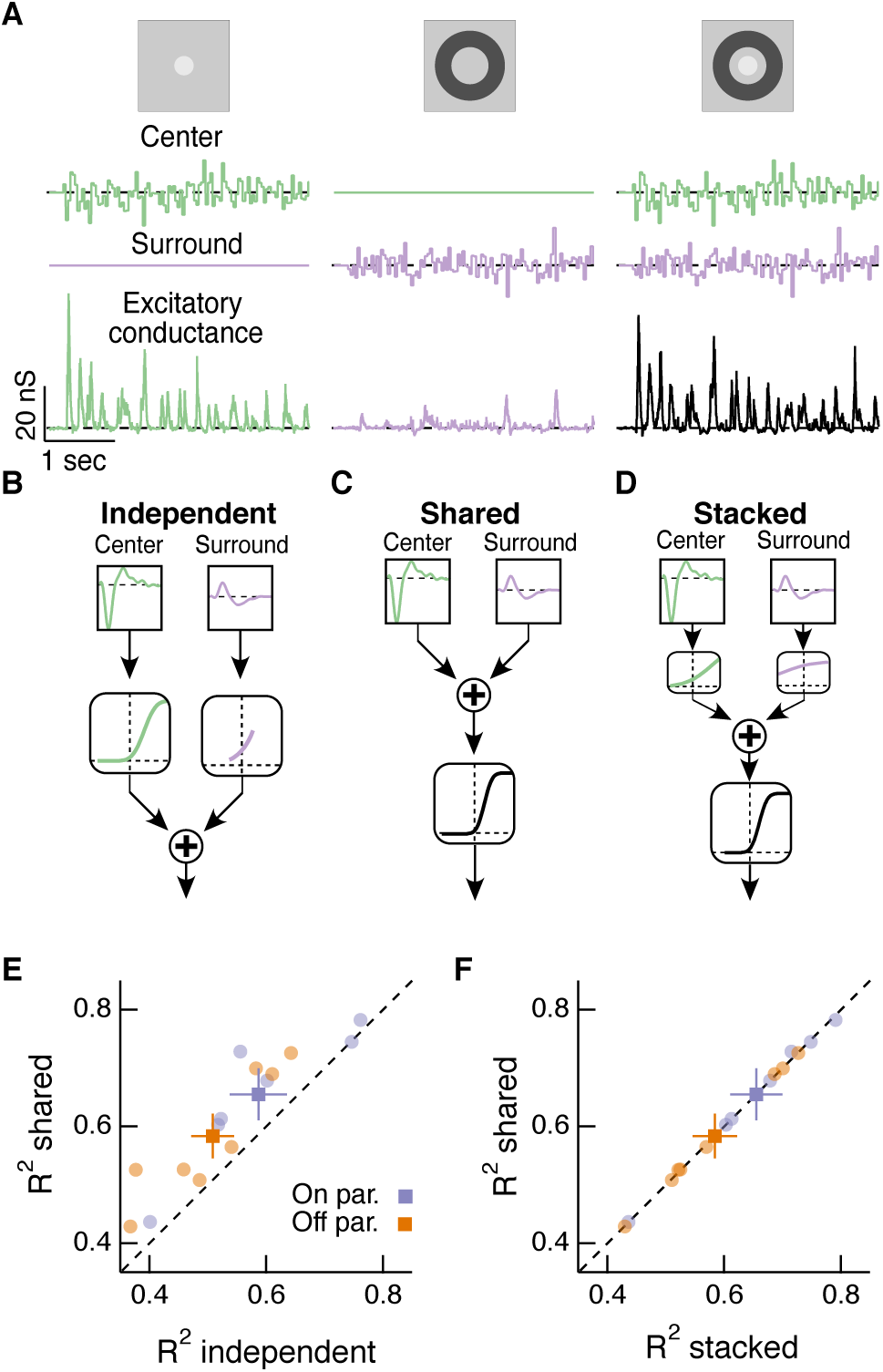
Linear-nonlinear cascade modeling supports an architecture where center and surround combine linearly before passing through a shared nonlinearity. (A) We presented Gaussian white noise to either the center (left), surround (middle) or center and surround simultaneously (right) while measuring excitatory synaptic current responses. Note that we’ve converted measured excitatory currents to an estimate of the excitatory conductance (in nS). Example traces are from a representative Off parasol RGC. (B) The independent model treats the filtered center and surround inputs with private nonlinear functions, and the outputs of these two nonlinearities are then summed to produce the excitatory conductance response. (C) The shared model integrates filtered center and surround inputs linearly, and this summed input is then passed through a single, shared nonlinearity. (D) The stacked model combines the independent and shared models by treating center and surround with independent nonlinearities before summation and treatment with a third, shared nonlinearity. (E) We tested the ability of each of these models to predict held-out responses to center-surround stimulation. (n = 7 On cells, *p* = 0.02; n = 8 Off cells, *p* = 0.002). (F) The fraction of explained variance was the same for the shared compared to the stacked model (*p >* 0.90 for both On and Off cells).

We constructed three models of how center and surround inputs combine to determine the cell’s excitatory conductance response when both RF regions are stimulated (see Methods for details). (1) In the “independent” model (Fig. 4B), inputs to the center and surround are filtered using their respective linear filters and then passed through separate nonlinear functions. The outputs of the two nonlinearities are then summed to give the response of the cell. (2) In the “shared” model (Fig. 4C), filtered center and surround inputs are summed linearly before passing through a single, shared nonlinear function. The output of this nonlinearity is the cell’s response. (3) The “stacked” model (Fig. 4D) combines models (1) and (2) by treating center and surround with private nonlinearities before summation and treatment with a third, shared nonlinearity. Representative predictions and measured responses are shown in Fig. S3. The shared model is a special case of the stacked model, where the upstream independent functions are linear. Similarly, the independent model is a special case of the stacked model, where the output function after summation is linear.

We fit the nonlinear functions in each model using a subset of simultaneous center-surround trials and used the remaining simultaneous trials to test how well each model could predict the cell’s response. The shared model outperformed the independent model (Fig. 4E). The shared model performed as well as the more complicated stacked model (Fig. 4F), despite the latter having many more free parameters (10 free parameters) than the shared model (5 free parameters). In addition, the private nonlinearities fit in the stacked model tended to be quite shallow and much nearer to linear than the sharply rectified shared nonlinearity (Fig. 4D). The modeling result in Fig. 4F supports the hypothesis that the dominant nonlinear interaction between center and surround is characterized by a shared nonlinearity, and that upstream of this nonlinearity center and surround interact approximately linearly.

The hypothesis in Fig. 1 suggests that the effective rectification experienced by each subunit will depend on surround activation. The experiments used for the modeling above allowed us to directly examine whether this is the case. To do this, we estimated center and surround activation by convolving center and surround filters with the appropriate stimuli. We then plotted the measured excitatory conductance against these estimates of center and surround activation; Fig. 5A shows an example for the same Off parasol RGC as Fig. 4A-D.

**Figure 5:**
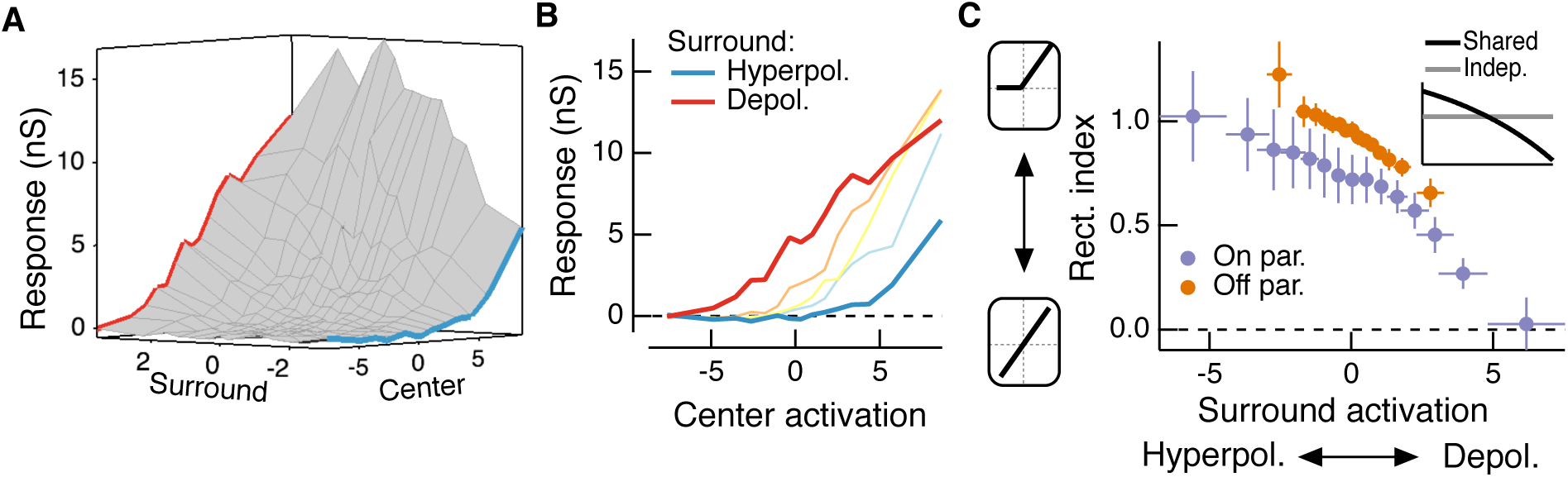
The RF surround changes the apparent rectification of inputs from the center. (A) Response surface showing the mean excitatory conductance response from an Off parasol RGC as a function of filtered inputs to both the center and surround (center or surround “activation”, i.e. their generator signals). (B) Sections through this surface at various levels of surround activation reveal that the shape of the nonlinear dependence of excitatory conductance on center activation changes as the surround is modulated. (C) To quantify this change in center rectification, we used a rectification index (see text), where values near 0 indicate a linear relationship between center activation and conductance response, and values near 1 indicate a sharply rectified relationship. Points are mean *±* S.E.M. (n = 7 On parasol cells and n = 8 Off parasol cells). Inset shows the relationship between rectification index and surround activation for a shared nonlinearity model (black curve) and an independent nonlinearity model (gray curve).

These joint response distributions show that the relationship between center activation and excitatory conductance depends on surround activation. When the surround is only weakly activated (near zero on “surround” axis in Fig. 5A), this nonlinear relationship is rectified (Fig. 5B). Rectification persists when the surround hyperpolarizes presynaptic bipolar cells (negative on “surround” axis in Fig. 5A, blue trace in Fig. 5A,B). But when the surround depolarizes bipolar cells (positive on “surround” axis in Fig. 5A), the relationship between center activation and excitatory conductance becomes more linear (i.e. less rectified; Fig. 5A,B, red trace). We quantified this change in center rectification with surround activation using a rectification index (RI, see Methods for calculation of this metric). A RI value of zero indicates a linear relationship between center activation and conductance response, whereas RI values near 1 indicate strong rectification (i.e. there is a large increase in response with positive center activation, but very little or no decrease in response with negative center activation). For both On and Off cells, rectification decreased as surround activation increased (Fig. 5C). In agreement with previous observations [47, 42], Off cells were more rectified than On cells. The inset to Fig. 5C shows the relationship between surround activation and RI for independent and shared nonlinearity models. When center and surround nonlinearities are independent, the rectification of the center does not depend on the activity of the surround (horizontal gray line in inset). The shared nonlinearity model, on the other hand, shows a decrease in rectification as the surround becomes more depolarizing, in qualitative agreement with the behavior of parasol RGCs.

### RF center and surround interact nonlinearly during naturalistic visual stimulation

To test whether inputs to the RF center and surround interact nonlinearly under naturalistic stimulus conditions, we used a visual stimulus designed to approximate natural primate viewing conditions based on the Database Of Visual Eye movementS (DOVES, [48, 49]). An example image and a corresponding eye movement trajectory is shown in Fig. 6A. We masked the stimulus to the RF center, surround or both regions. Fig. 6B shows spike responses of an example Off parasol RGC to three movie stimuli: stimulation of the center alone (Fig. 6B, green), stimulation of the surround alone (Fig. 6B, purple), or simultaneous stimulation of both the center and surround (Fig. 6B, black). Responses to isolated center or surround stimuli are shown in Fig. 6C for average spike responses (Fig. 6C, top) and excitatory synaptic inputs (Fig. 6C, bottom).

**Figure 6:**
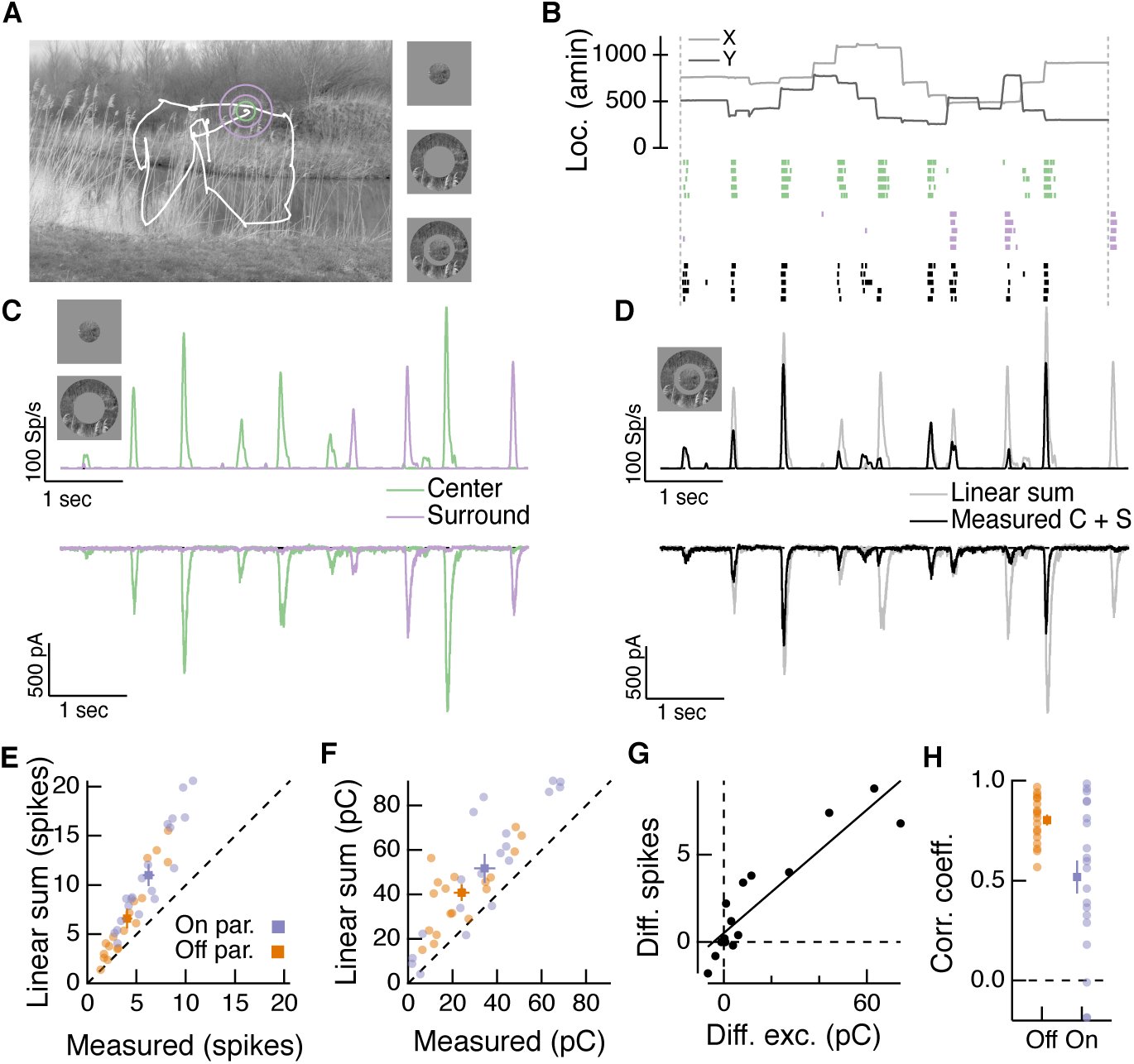
Natural movie stimuli elicit nonlinear interactions between the RF center and surround. (A) Natural image and associated eye movement trajectory from [48]. Right: example movie frames showing isolated center (top), surround (middle), and center-surround stimuli (bottom). (B) Rasters show example Off parasol RGC spike responses to these three movie stimuli. Top shows eye movement position. (C) Spike output (top) and excitatory synaptic input (bottom) responses to isolated center and surround stimuli. (D) Spike and excitatory synaptic input responses to the center-surround stimulus. Gray trace shows the linear sum of isolated center and surround responses. (E) We compared the spike count in response to the center-surround stimulus to the linear sum of isolated center and surround response magnitudes. Each point is a different natural movie. Center and surround sum sub-linearly (On parasol RGCs: n = 20 natural movies across 8 cells, *p <* 3 *×* 10^−7^; Off parasol RGCs: n = 18 natural movies across 7 cells, *p <* 2 *×* 10^−4^). (F) Same as (E) but for excitatory charge transfer responses (On parasol RGCs: *p <* 3 *×* 10^−5^; Off parasol RGCs: *p* = 8 *×* 10^−6^). (G) For the example in (A-D), the difference between measured and linearly-summed spike responses was correlated with differences in excitatory synaptic inputs (r = 0.91). (H) Population data for the analysis in (G).

To determine whether center and surround signals interact nonlinearly, we compared the linear sum of center and surround responses (Fig. 6D, gray traces) to the measured response to simultaneous stimulation of both the center and surround (Fig. 6D, black traces). For both spike and excitatory current responses, the measured center-surround response was smaller than the linear sum of the two responses measured independently. Thus RF center and surround interact nonlinearly. As in Figs. 4 & 5, because this interaction is present in the excitatory synaptic input, it is not due to nonlinearities in synaptic integration or spike generation in the ganglion cell.

Sublinear interactions between center and surround inputs held across cells and fixations for both spike output (Fig. 6E) and excitatory synaptic input (Fig. 6F). For each cell, the difference between the linear sum of center and surround inputs and the measured simultaneous response was similar for excitatory inputs and spike outputs (Fig. 6G,H). This is consistent with the interpretation that the nonlinear interaction seen at the level of spike output is inherited from the excitatory synaptic inputs. Correlations between sublinear interactions in spike output and excitatory synaptic input were stronger for Off compared to On parasol RGCs (Fig. 6H). This may be because inhibitory input has a strong impact on On but not Off parasol RGC spike responses to natural stimuli [42]. Taken together, these observations demonstrate that the nonlinear center-surround interactions characterized in Figures 2-5 are prominent for naturalistic visual inputs.

### Natural spatial correlations promote nonlinear center-surround interactions

How do nonlinear center-surround interactions depend on stimulus statistics, especially those that characterize natural scenes? Naturalistic center and surround stimuli tended to elicit responses at non-overlapping periods of the stimulus (Fig. 6). This is consistent with the spatial correlations in intensity that characterize natural images [4] and the antagonistic nature of the surround—e.g. an Off-center parasol RGC would be depolarized by negative contrast in the RF center and hyperpolarized by negative contrast in the surround. We tested the effect of spatial correlations on nonlinear center-surround interactions using a synthetic visual stimulus inspired by our natural movie stimuli.

This stimulus consisted of a uniform disc in the center and a uniform annulus in the surround. The intensity of each region was sampled from a natural image (Fig. 7A,B) and presented to either the center, surround, or center and surround simultaneously. To choose the center and surround intensities, we computed the mean intensity within the disc and annulus for randomly chosen image locations. The intensity correlations characteristic of natural scenes were evident when we plotted the center intensity against the corresponding surround intensity (Fig. 7C, left, “Control”). Shuffling the surround intensities relative to those of the center eliminated those spatial correlations, while maintaining the same marginal distributions (Fig. 7C, right, “Shuffled”).

**Figure 7:**
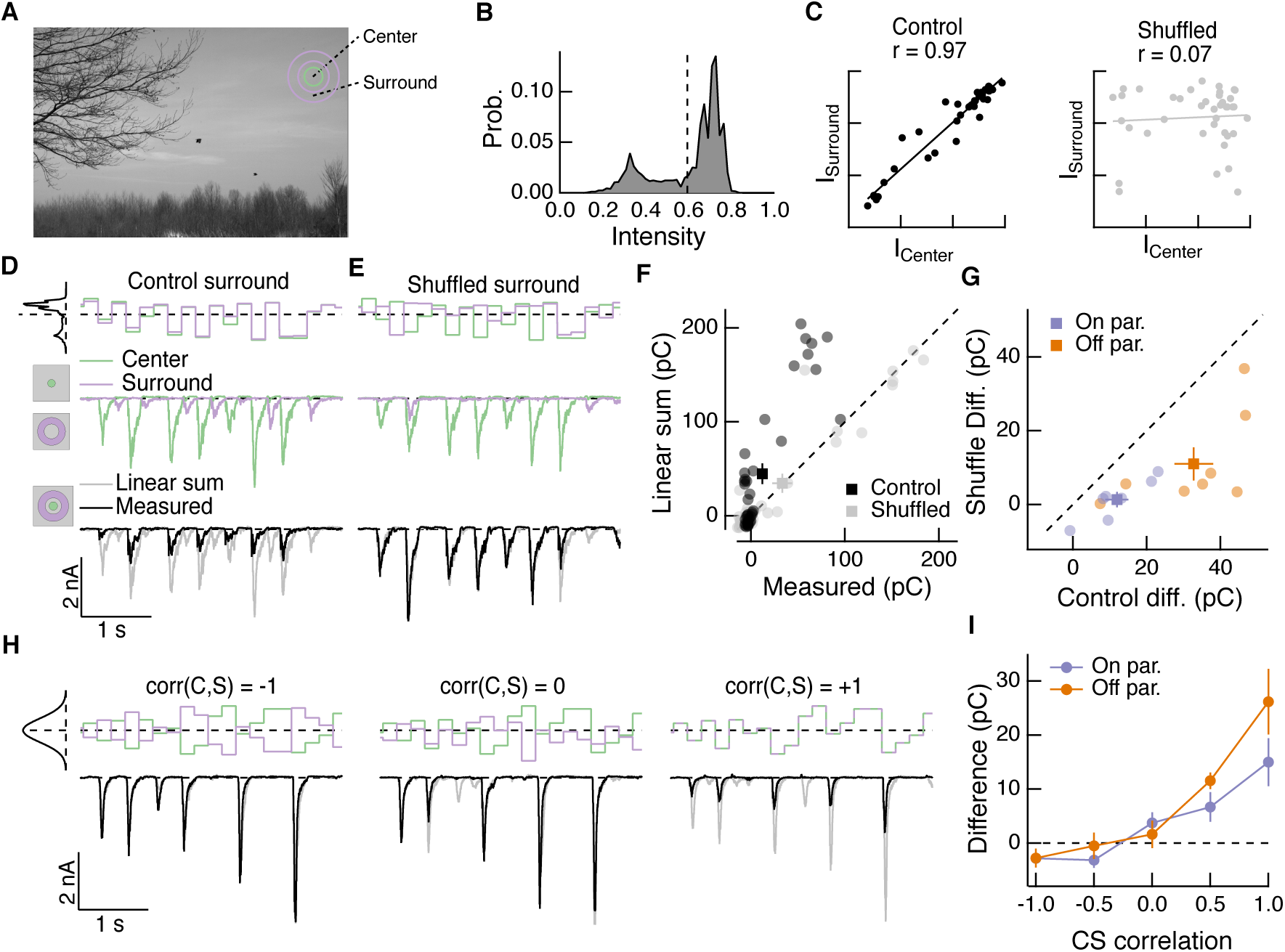
Spatial correlations in natural scenes promote nonlinear center-surround interactions. (A) Example image [49] used to construct natural intensity stimuli. (B) Intensity histogram from the image in (A). Dashed vertical line indicates the mean intensity, which was used as the mean gray level in experiments that follow. (C) Center and surround intensity values for 40 image patches from the image in (A). (D) Example stimuli (top) and Off parasol RGC excitatory current responses to isolated center and surround (middle) and center-surround (bottom) stimulation. Gray trace in bottom shows linear sum of isolated center and surround responses. (E) Same as (D) for shuffled surround intensities.(F) The response magnitude (charge transfer) of each fixation is plotted for measured center-surround and linearly-summed center and surround responses. Circles show mean responses for each fixation, squares show mean *±* S.E.M. across all fixation responses in this example cell. (G) Population data showing the mean difference between responses to the center-surround stimulus and the linearly-summed response. Circles show average differences for each cell tested, and squares show population mean *±* S.E.M (n = 7 On parasol RGCs, *p <* 4 *×* 10^−4^; n = 8 Off parasol RGCs, *p <* 2 *×* 10^−3^). (H) We generated white noise center-surround stimuli that had variable center-surround correlations but constant marginal distributions. Shown are example excitatory current responses in an Off parasol RGC. Black traces show the measured center-surround stimulus response and gray traces show the linear sum of center and surround responses. (I) Population data from the experiments in (H) showing that nonlinear center-surround interactions depend on the correlation between center and surround inputs (n = 8 On parasol RGCs; n = 8 Off parasol RGCs).

We presented a uniform-intensity disc to the center and/or annulus to the surround, updating the intensity of each region every 200 msec, which is consistent with typical human fixation periods [48]. When spatial correlations were intact, inputs to the center and surround combined sub-linearly in the excitatory synaptic input to the cell (Fig. 7D), as they did in the full natural movie responses (Fig. 6). When we shuffled the surround intensities relative to the center, nonlinear center-surround interactions were much weaker (Fig. 7E,F). This was true for excitatory synaptic inputs to both On and Off parasol RGCs (Fig. 7G).

To further probe the impact of intensity correlations on center-surround interactions, we generated Gaussian random noise stimuli that updated with a 200 msec period. For this stimulus, a single random intensity fills the entire center disc and a different, random intensity fills the entire surround annulus. This noise stimulus had a tunable degree of correlation between center and surround intensity, ranging from −1 (perfectly anti-correlated) to 0 (uncorrelated) to +1 (perfectly correlated, i.e. modulated in unison). When noise stimuli in the center and surround were negatively correlated, inputs to the center and surround summed linearly or very nearly so (Fig. 7H, left). As the center-surround intensity correlations increased, sublinear interactions became more obvious (Fig. 7H, middle and right). Strongly positively-correlated noise stimuli induced center-surround interactions that resembled those seen using naturally-correlated luminance stimuli (Fig. 7H, right). This dependence of nonlinear center-surround interactions on center-surround intensity correlations was present in both On and Off parasol RGCs (Fig. 7I).

### A luminance-matched surround promotes spatial contrast sensitivity in the center

The experiments described above show the surround signals, rather than only interacting with the fully formed center signal, can alter how the center integrates over space. Two aspects of these results stand out: (1) nonlinear spatial integration is maximized when center and surround experience similar mean luminances (Figures 2, 3), and (2) natural stimuli elicit strong nonlinear center-surround interactions due to positive correlations between center and surround (Figures 6, 7). These observations led to the hypothesis, tested below, that surround activation can make spatial integration in the RF center relatively insensitive to mean luminance.

Visual stimuli, such as the change in input encountered after a saccade, typically include changes in mean luminance and spatial contrast. Such stimuli will activate both linear and nonlinear response components, and these may not interact in a straightforward manner. To make this more concrete, consider a single Off bipolar subunit. At rest, the bipolar synapse is in a sharply rectified state; any visual inputs that alter the bipolar cell voltage will alter the synaptic operating point and hence change the degree of rectification applied to a subsequent input. A decrease in mean luminance over the RF center will depolarize the synapse associated with each RF subunit, shifting each into a locally linear state (Fig. 8A, top), and decreasing spatial contrast sensitivity. When such a stimulus is paired with a luminance-matched surround stimulus, the antagonistic surround may shift each subunit back into a locally rectified state (Fig. 8A, bottom, blue arrow), restoring spatial contrast sensitivity.

**Figure 8:**
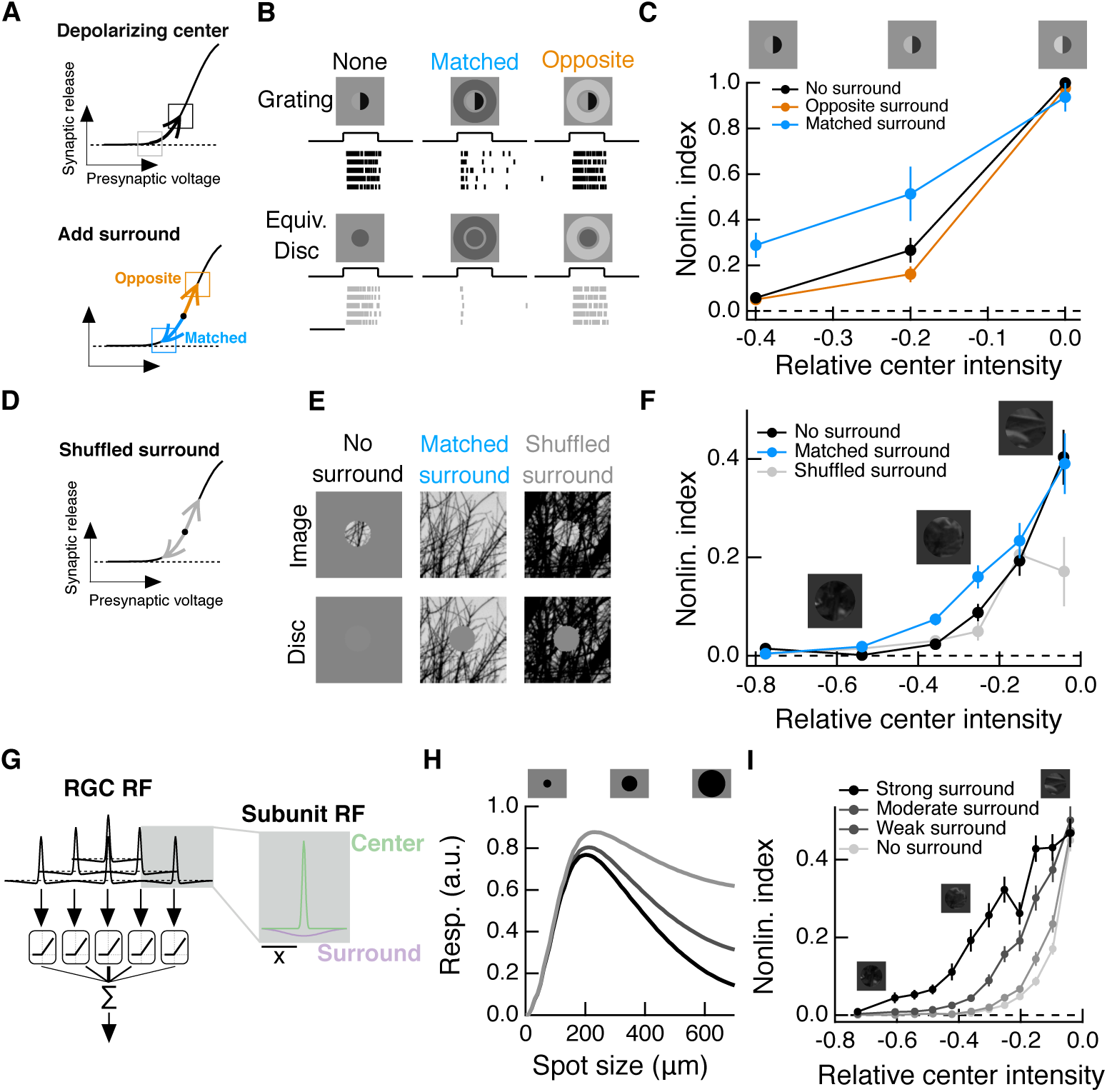
Intensity correlations across space promote nonlinear spatial integration in the RF center. (A) Schematic showing the hypothesized interaction between center and surround inputs on local subunit rectification. A depolarizing input to the center may push the synapse into a locally-linear state. A simultaneous surround input that is matched in luminance (blue arrow) can hyperpolarize the synaptic terminal and bring the synapse back into a rectified state, whereas a poorly matched surround will not (orange arrow). (B) During Off parasol spike recordings, we presented split-field grating stimuli to the RF center under three surround conditions. (C) Summary data showing the population mean *±*S.E.M. NLI (see text) as a function of the mean intensity (relative to the background) of the center grating (n = 8 Off parasol RGCs). (D-F) We presented natural image patches and their linear equivalent disc stimuli to measure the NLI under three surround conditions: no surround, a matched surround image, and a shuffled surround image. (G) Schematic of a nonlinear subunit RF model. Each subunit has a difference-of-Gaussians spatial receptive field. The output of each subunit is passed through a private, rectifying output nonlinearity. Subunit outputs are then summed over visual space to yield the modeled RGC response. (H,I) We changed the strength of the subunit surround to model RGCs with three different surround strengths: a weak surround (light gray trace), an intermediate-strength surround (gray trace), and a strong surround (black trace). We presented this RF model with the natural image/disc stimuli shown in (E) and, following that analysis, measured the NLI as a function of the mean intensity of the image in the RF center.

We tested this prediction using modified grating stimuli with nonzero mean luminance. For each grating stimulus, we also presented a corresponding linear equivalent disc stimulus, which has the same mean luminance as the grating, but lacks any spatial contrast. The degree to which a cell’s response to these two stimuli differs is a measure of the sensitivity of the cell to spatial contrast, or, equivalently, the strength of nonlinear spatial integration.

We presented these stimuli to Off parasol RGCs while measuring spike responses. A grating with a dark mean luminance signal produced a similar response compared to a linear equivalent stimulus (Fig. 8B, left), indicating that the cell is insensitive to the spatial contrast present in this grating stimulus. Compare this to these cells’ highly nonlinear responses to zero-mean grating stimuli (e.g. Fig. 3). When a dark mean grating is paired with a luminance-matched surround, however, the grating again produces a much stronger response than its linear equivalent stimulus (Fig. 8B, middle), which is consistent with the restoration of spatial contrast sensitivity via the mechanism in Fig. 8A. Pairing the grating stimulus with poorly-matched surrounds does not restore spatial contrast sensitivity (Fig. 8B, right), indicating that this is not a general consequence of surround activation.

To quantify the spatial contrast sensitivity in these experiments, we used a metric we call the nonlinearity index (NLI, See equation 1 and [42]):

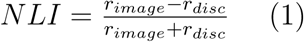

This measure normalizes responses within each surround condition. A positive NLI indicates that the cell responds more strongly to a grating stimulus than to its linear equivalent disc stimulus, and is thus sensitive to spatial contrast. A NLI near zero indicates that the cell’s response is mostly determined by the mean luminance component of the stimulus, and not the spatial contrast. When the mean intensity of the grating was equal to the background intensity, the NLI was maximal for all surround conditions. As the center intensity decreased (moving towards the left on Fig. 8C), the NLI decreased along with it, but this drop was less pronounced under the matched surround condition. Hence matched surround activation decreased the sensitivity of nonlinear spatial integration to changes in mean luminance.

### Intensity correlations in natural images promote nonlinear spatial integration

The results presented thus far show that systematically varying the input to the surround relative to the center can alter sensitivity to spatial contrast in both artificial and natural image stimuli (Figures 2, 3, 8A-C). What is not clear from these experiments is how much this effect is present during the course of more naturalistic activation of the RF surround. The intensity correlations present in natural images (e.g. Fig. 7C) should ensure that the mean intensity difference between the RF center and surround is often near zero. Because spatial contrast sensitivity was maximized by small differences between mean center and surround intensity (Fig. 2), we hypothesized that full natural image stimulation of the RF surround would increase spatial contrast sensitivity in the RF center compared to stimulation of the RF center alone.

To test this hypothesis, we presented natural image patches (and their corresponding linear equivalent disc stimuli) to the RF center while pairing each with three distinct surround conditions: no surround stimulation (Fig. 8E); the naturally-occurring surround present in the rest of the natural image patch (“Matched surround”), and, a randomly-selected surround from a different region of the same full natural scene (“Shuffled surround”). We recorded Off parasol spike responses to these six stimuli for each of 20-40 randomly-selected image patches from a single natural scene. For each image patch, we computed the NLI (Equation 1) for responses measured in each of the three surround conditions. We then compared the NLIs to the mean intensity of the image in the RF center (I) relative to the background intensity (B), i.e. Relative center intensity = (*I - B*)*/B*. As in the modified gratings experiments (Fig. 8C), a darker mean luminance signal was associated with a decrease in spatial contrast sensitivity (Fig. 8F, black curve). This drop-off in spatial contrast sensitivity was less pronounced with the naturally-occurring surround, (Fig. 8F, blue curve). Specifically, sensitivity to fine spatial structure was 2-3 times greater in the presence of a matched surround than without a surround. Randomly-selected surrounds did not enhance spatial contrast sensitivity in this way but instead altered the NLI in a manner predicted by the experiments in Fig. 2 (see Fig. S4).

These observations are consistent with the idea that natural images provide inputs to the surround that can preserve the spatial contrast sensitivity of the RF center compared to center inputs alone. Key to this relative invariance of contrast sensitivity are the ability of the surround to control the degree of rectification of the bipolar subunits that comprise the RF center and the strong positive correlations between center and surround inputs created by natural images.

To explore the relationship between naturalistic surround activation and spatial contrast sensitivity in a manner not possible in our experiments, we constructed a simple spatial RF model composed of nonlinear, center-surround subunits (Fig. 8G; see [38] and Methods). Following the analysis used for the data in Fig. 8F, we computed the mean NLI for the model as a function of the relative center intensity for a surround-free stimulus (Fig. 8I, “no surround”) and for the naturally-occurring surround stimulus (Fig. 8I, “moderate surround”). As in the Off parasol spike data, the inclusion of the naturally-occurring surround extended spatial contrast sensitivity in the face of stronger local luminance signals. We repeated the same analysis for versions of the RF model with both a weaker and a stronger surround. A stronger RF surround is associated with greater spatial contrast sensitivity, especially for images that contain a strong local luminance signal. Similar results were seen for a spatiotemporal RF model that includes temporal filters measured in the experiments shown in Fig. 4 (Fig. S5).

## Discussion

Here we have shown that visual inputs to the RF surround can modulate the spatial integration properties of the center. Functionally, this means that inputs to the surround can change how the RF center integrates inputs across space, including in natural scenes. During the course of natural viewing conditions, the local luminance experienced by a RGC can vary dramatically from fixation to fixation [50]. Because sensitivity to small scale (sub-RF center) spatial contrast relies critically on the rectification of subunit outputs, these changes in luminance could prevent a RGC that responds nonlinearly to grating stimuli from similarly detecting spatial contrast present in natural images. The incorporation of an antagonistic surround before formation of the nonlinear center ensures that this effect of the mean intensity signal is mitigated to some degree for natural scenes and other stimuli with strong correlations in intensity. The degree to which spatial contrast sensitivity is preserved will depend on the strength of the surround, and hence both cell type and properties of the visual environment that can alter surround strength like ambient luminance levels [51]. For those (relatively rare) regions of an image where the center and surround luminance signals are very different, a nonlinear RGC will lose much of its small-scale spatial contrast sensitivity and instead encode input with a spatial scale dictated by the classical center-surround RF. This could happen in a region of a scene that contains structure like an edge or a boundary between two large objects.

Past work in parasol RGCs as well as other RGC types has shown that the inclusion of nonlinear subunits in the RF center is important to capture responses to spatially structured stimuli [52, 44, 53, 54, 39], including natural scenes [42]. The results shown here rely on nonlinear subunits in the RF center having a center-surround RF organization. This is consistent with previous observations that the surround is present in diffuse bipolar cells in primate retina [33], and with past models of cat Y-cell RGCs [38]. The work presented here indicates that it will be important to incorporate the impact of the surround on subunits within the receptive field center to capture RGC center-surround interactions and spatial contrast sensitivity.

For the sake of concreteness, and based on past work in mouse retina [41], we have assumed that the nonlinear relationship between bipolar cell input and excitatory input to the RGC arises at the bipolar cell to RGC synapse, but we note that some contribution from nonlinearities earlier in the bipolar cell is likely as well. For example, active conductances in the axons of diffuse bipolar cells would also be expected to shape the nonlinear relationship between bipolar cell input and synaptic release [55]. Modulation of the bipolar cell synaptic operating point can result from changes in mean luminance [41] or from genetic or pharmacological manipulations that change the resting membrane potential of AII amacrine cells [56]. The principle here is similar except in this case the synaptic operating point is being dynamically modulated with surround activation.

Similar RF subregion interactions are present in other visual neurons, as well. For example, activation of suppressive subunits in V1 RFs can change the shape of the nonlinear relationship between excitatory subunit activation and spike response [57]. V1 surrounds are recruited when inputs to the surround are similar to those in the center [58]. The result is that the V1 surround is strongest in homogenous visual contexts. Here we see a similar contextual effect of retinal surrounds, albeit with a sensitivity to lower-level statistical features. The surround has the greatest impact on RGC responses when visual stimuli contain luminance correlations (Figs. 6,7).

The circuit mechanisms that give rise to the observed center-surround interactions and modulation of spatial contrast sensitivity, namely nonlinear synaptic transfer functions and convergence of parallel neural pathways, are ubiquitous in neural circuits beyond the retina. This work supports the broader notion that complex neural computations need not rely on exotic circuitry. Rather, interactions of known circuit elements can produce a variety of computations under the right stimulus conditions by producing modest changes in signaling that enhance or mitigate the impact of nonlinear mechanisms such as the bipolar output synapse. This highlights the importance of exploring how known neural circuit mechanisms interact under diverse stimulus conditions, including during naturalistic stimulation.

## Methods

### Tissue preparation

We obtained retinal tissue from Macaque monkeys (*M. nemestrina, M. mulatta*, or *M. fascicularis*) via the tissue distribution program at the Washington National Primate Research Center. All procedures were approved by the Institutional Animal Care and Use Committee at the University of Washington. Dissection procedures have been described previously [59, 42]. After enucleation, the eye was hemisected and the vitreous humor was removed mechanically, sometimes assisted by treatment with human plasmin (*∼*50 *µ*g/mL, Sigma or Haematologic Technologies Inc.). Retina was dark adapted for *∼*1 hr, and all subsequent procedures were performed under infrared light using night-vision goggles. The retina and pigment epithelium were separated from the sclera and stored in oxygenated (95%O_2_/5%CO_2_) Ames bicarbonate solution (Sigma) in a light-tight container. Retinal mounts were removed from the pigment epithelium and laid photoreceptor-side down onto a poly-D-lysine coated coverslip (BD biosciences). During experiments, retinal tissue was perfused at 7-9 mL/min with Ames solution at *∼*32°C.

### Patch recordings

Electrophysiological recordings were performed using a Multiclamp 700B amplifier (Molecular Devices). Spike responses were measured using extracellular or loose-patch recordings with an Ames-filled pipette. For voltage clamp recordings, we used low-resistance pipettes (tip resistance *∼*1.5-4 MΩ) filled with a Cs-based internal solution (containing, in mM: 105 CsCH3SO3, 10 TEA-Cl, 20 HEPES, 10 EGTA, 5 Mg-ATP, 0.5 Tris-GTP, and 2 QX-314, pH 7.3, *∼*280 mOsm). We compensated for access resistance (*∼*4-8 MΩ) online by 75%. Reported voltages have been corrected for an approximately −10 mV liquid junction potential. To measure excitatory synaptic inputs in voltage clamp recordings, we held the cell at the expected reversal potential for inhibitory inputs. This was typically around −60 mV, but was adjusted for each cell by delivering light steps at holding potentials near this value until the inhibitory response was eliminated.

### Cell identification and selection

We identified On and Off parasol RGCs under infrared illumination based on soma size and morphology as well as characteristic spike responses to light steps centered on the cell. The overall health and sensitivity of the retina was confirmed by delivering a uniform, 5% contrast, 4 Hz modulated stimulus, which produces a robust spike response (at least a few spikes per cycle) in On parasol RGCs in sufficiently sensitive tissue. Sensitivity was continuously monitored (typically before each recording) in this way.

### Visual stimulation

Stimuli were presented and data acquired using custom written stimulation and acquisition software packages Stage (stage-vss.github.io) and Symphony (symphony-das.github.io). Lab-wide acquisition packages can be found at https://github.com/Rieke-Lab/riekelab-package and protocols used in this study can be found at https://github.com/Rieke-Lab/turner-package.

Visual stimuli were presented with 60Hz frame rates on an OLED microdisplay monitor (eMagin, Bellevue, WA) focused onto the photoreceptors. Monitor outputs were linearized by gamma correction. Stimuli were calibrated using monitor power outputs, the spectral content of the monitor, macaque photoreceptor spectral sensitivity [60], and a collecting area of 0.37 *µm*^2^ for cones [61] and 1 *µm*^2^ for rods. Unless otherwise noted, mean light levels produced *∼*9,000 isomerizations (R*)/M or L-cone/s, *∼*2,000 R*/S-cone/s and *∼*18,000 R*/rod/s.

Before every parasol RGC recording, we found the center of the cell’s RF using a split-field contrast reversing grating stimulus at 4 Hz and 90% contrast. To do this, we translated the grating until the two F2 response cycles were balanced (i.e. we minimized the F1 while maximizing the F2 component of the response). We performed this search in both the horizontal and vertical dimensions. In experiments where we specifically targeted visual stimuli to the RF center or surround, we measured each cell’s area-summation curve and fit, online, to these data a circular difference-of-Gaussians RF model in radial coordinates. This model is described in equation 2 below.

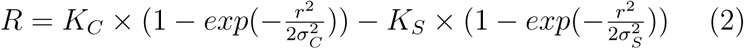

Where *R* is the response of the cell, *r* is the spot radius, and four free parameters describe the shape of the RF, two for each center and surround: *K* describes the amplitude scaling of each component and *σ* describes the size. We then chose a boundary for the center that would minimally activate the surround while still filling much of the center and likewise for the surround (e.g. see Fig. S1). These boundaries were used to generate appropriate masks and apertures to target each RF subregion.

### Natural visual stimuli

Naturalistic movie stimuli (Fig. 6) were generated using data from the DOVES database [48] http://live.ece.utexas.edu/research/doves/. The images in the DOVES database were selected from the van Hateren natural image database [49], and we used the original van Hateren images instead of the images included in the DOVES database. This is because the DOVES images are cropped, and using the original, larger images allowed for a wider selection of eye movement trajectories to be used. To select data from the DOVES database to use in experiments, we ensured that the eye trajectory never extended close enough to the boundaries of the image that our presented frames would extend beyond the boundaries of the image.

For natural movie (Fig. 6), luminance (Fig. 7), and image stimuli (Fig. 2,8), we scaled each image such that the brightest pixel in the image was assigned an intensity value of 1 (maximum monitor intensity). The mean gray level of the monitor was set to the mean pixel intensity over the entire image. This mean gray level was used in masked/apertured regions of the frame as well as the blank screen between trials. Natural movies were presented at a spatial scale of 1 arcmin/pixel, which is equal to 3.3 *µm*/pixel on the monkey retina. Natural images, from the van Hateren database, were presented at a scale of 6.6 *µm*/pixel on the retina.

In the experiments in Fig. 7, we updated the intensity of a disc (annulus) in the center (surround) every 200 msec, which is consistent with typical human fixation periods during free-viewing [48], but on the rapid side of the distribution (for efficient data collection). To compute natural intensity stimuli for the center and surround, we selected many random patches from a natural image and measured the mean intensity within a circle of diameter 200 *µm* (for the RF center) and within an annulus with inner and outer diameters 200 and 600 *µm*, respectively (for the RF surround).

For the natural image experiments in Fig. 2 we used a natural image patch selection method that has been described previously [42]. Briefly, we used a nonlinear subunit model developed in to rank image patches based on their degree of response nonlinearity (i.e. how differently they drove a spatially linear compared to a spatially nonlinear RF model). We then sub-sampled the entire distribution of image patches in order to present an approximately uniform distribution of images that ranged from very nonlinear (i.e. high spatial contrast) to very linear (i.e. low spatial contrast). This ensured that we could efficiently explore the full range of spatial contrasts present in natural images within a single experiment. For the natural image experiments and modeling in Fig. 8, we did not use this biased sampling method, instead selecting image patches randomly from within a natural image.

### Data analysis and modeling

Data analysis was performed using custom written scripts in MATLAB (Mathworks). The code used to analyze the data in this study, perform the computational modeling, and generate the figures presented can be found at https://github.com/mhturner/CenterSurroundInteractions, and more general analysis code used in this study can be found at https://github.com/mhturner/MHT-analysis. Throughout the study, reported p-values were computed using t-tests.

The center-surround models in Fig. 4 were constructed as linear-nonlinear cascade models. Using trials where only the center or surround was stimulated in isolation, we used reverse correlation between the cell’s response and the noise stimulus to compute the linear filter for both the center and the surround. Each of the three models presented in Fig. 4 took as inputs the filtered center and surround noise stimuli.

For each of the nonlinearities in the models, we parameterized a smooth curve based on a cumulative Gaussian function to fit to the mean response as a function of generator signal (see [62]). Here, *C*() is the cumulative normal distribution (*normcdf* in MATLAB).

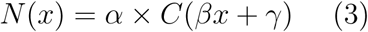

Equation 3 above shows the general form of this nonlinear function, which is determined by three free parameters: *γ* is an offset along the horizontal axis, *β* determines the sensitivity or slope of the contrast response function, and *α* is a scale factor which determines the maximal response. Equations for the three models are described in more detail below.

1. The **Independent model** is described by seven free parameters. Three each for each of the independent nonlinearities of the form described in equation (3) above, and one, *∊*, which sets the vertical offset of the output response. In this and the following equations, *h*_*c*_ (*h*_*s*_) is the linear filter for the RF center (surround), *s*_*c*_ (*s*_*s*_) is the stimulus in the center (surround) and the *\** operator indicates a linear convolution.

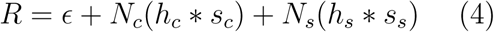
2. The **Shared model** is described by five free parameters. Three for the single nonlinearity (equation 3), a vertical offset of the response (*∊*), and a scale factor, *a*, which sets the relative weight of the center compared to the surround.

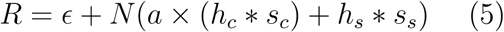
3. The **Stacked model** is described by ten free parameters. Three for each of the three nonlinearities (equation 3) and a single vertical offset, *∊*.

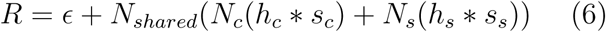

To fit and test these models, we collected *∼*30 trials total, ten each of center, surround, and center-surround conditions. We tested the models’ predictive power against a single center-surround trial, fitting the model with the remaining trials. We repeated this for every trial, holding each out from the model fitting for testing purposes. The reported r-squared values are the average over these fitting iterations.

For the image computable model in Fig. 8, we converted image patches to Weber contrast (*C*) by normalizing each pixel intensity (*I*) to the mean over the entire natural image (*B*), i.e.

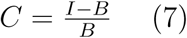

The models in Fig. 8 and S5 are of an Off-center RF, so contrast-converted images were multiplied by −1. For both expanding spots stimuli and natural images in the spatiotemporal model, stimuli were turned on for 200 msec from a mean-gray level, as in the experiments. To compute the spatial activation of each subunit, we first convolved an image stimulus with one of two spatial filters: a Gaussian representing the subunit center, or a Gaussian representing the subunit surround. The parameters describing the RF sizes were not fit to cells, but selected to be representative of RF center, surround, and subunit sizes typical of Off parasol RGCs at the retinal eccentricities used for these experiments. Subunits were placed on a square grid in visual space. Each subunit’s RF sub-field activation (*a*_*c*_ for center activation, *a*_*s*_ for surround activation) was computed by sampling the appropriate convolved image at that subunit’s location. For the spatiotemporal model in Fig. S5, the “voltage” signal of each subunit (*V*_*i*_, i.e. the pre-nonlinearity activation), as a function of time is given by equation 8,

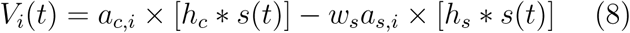

where *h*_*c*_ and *h*_*s*_ are the temporal linear filters for center and surround, respectively. The filters we used were Off parasol excitatory current population means from the center-surround white noise experiments in Fig. 4. ω_*s*_ is the weight of the subunit surround, relative to the center, and *s*(*t*) is the stimulus waveform (in this case, a 200 msec step from zero). Again, the *\** operator indicates a linear convolution. The final step in computing the spatiotemporal model’s response is to treat each subunit voltage, *V*_*i*_(*t*), with a rectifying nonlinear function and sum over all subunit outputs. This is given by equation 9,

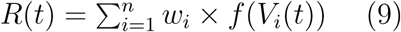

where ω_*i*_ is the weighting of each subunit, defined by a circular Gaussian function over the RF center; the nonlinear transform applied to each subunit activation, *f*, was a simple threshold-linear function. Examples of the time-varying model responses, *R*(*t*), can be seen in Fig. S5B. To compute the responses in Fig. S5C,D we integrated time varying responses over the duration of the 200 msec step (as in the analyses used in our experiments).

For the simpler spatial RF model (with no temporal component) in Fig. 8, the pre-nonlinearity Voltage of each subunit and response of the model RF is given by equations 10 and 11 below,

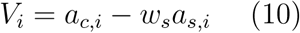

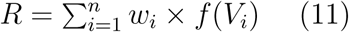

The RF model is described by four fixed parameters: the size of the Gaussian subunit center, (*σ* = 10 *µm* for the results shown here), the size of the subunit surround (*σ* = 150 *µm*), the size of the RF center, used to weight subunit outputs (ω_*i*_ in equation 9 above, *σ* = 40 *µm*), and the strength of the subunit surround, relative to the strength of the subunit center (0.5 for the “Weak surround” model, 1.0 for the “Moderate surround,” and 1.5 for the “Strong surround”).

The rectification index (RI) used in Fig. 5 is given by equation 12 below,

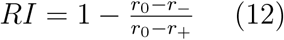

where *r*_0_ is the mean conductance response to zero center activation, *r*_*-*_ is the response to maximal center deactivation (i.e. negative center activation), and *r*_+_ is the response to maximal center activation.

## Author contributions

M.H.T., G.W.S. and F.R. conceived the project; M.H.T. and F.R. designed and conducted experiments; M.H.T. performed the analysis and modeling; M.H.T. and F.R. wrote the manuscript with input from G.W.S.

## Acknowledgments

We thank Shellee Cunnington and Mark Cafaro for excellent technical support. Tissue was provided by the Tissue Distribution Program at the Washington National Primate Research Center (WaNPRC), and we are grateful for assistance from the WaNPRC staff, especially Chris English and Drew May. Raunak Sinha and Mike Manookin assisted in tissue preparation. We thank Nora Brackbill, Jon Cafaro, Mike Manookin, Luis Gonzalo Sánchez Giraldo and Odelia Schwartz for helpful feedback on an earlier version of this manuscript. This work was supported by NIH grants F31-EY026288 (M.H.T.) and EY11850 (F.R.), and the Howard Hughes Medical Institute (F.R.).

## Supplementary information

**Figure S1:**
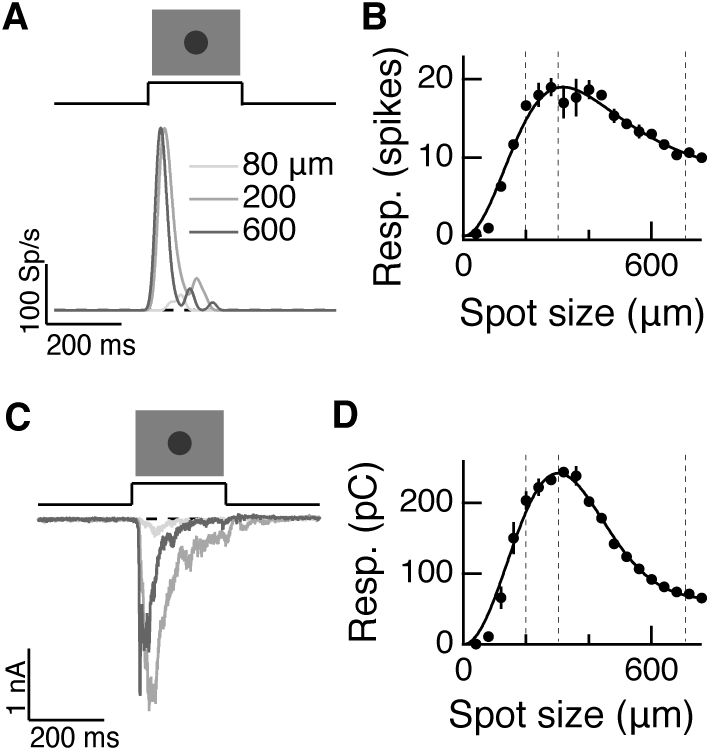
Measuring linear center-surround structure in parasol RGCs; To accompany Fig. 2. (A) Off parasol RGC spike responses to expanding spots stimuli. (B) Area-summation curve from the cell in (A), points show mean *±* S.E.M., smooth curve is a difference-of-Gaussians fit. (C-D) Same as (A-B) for excitatory current responses.

**Figure S2:**
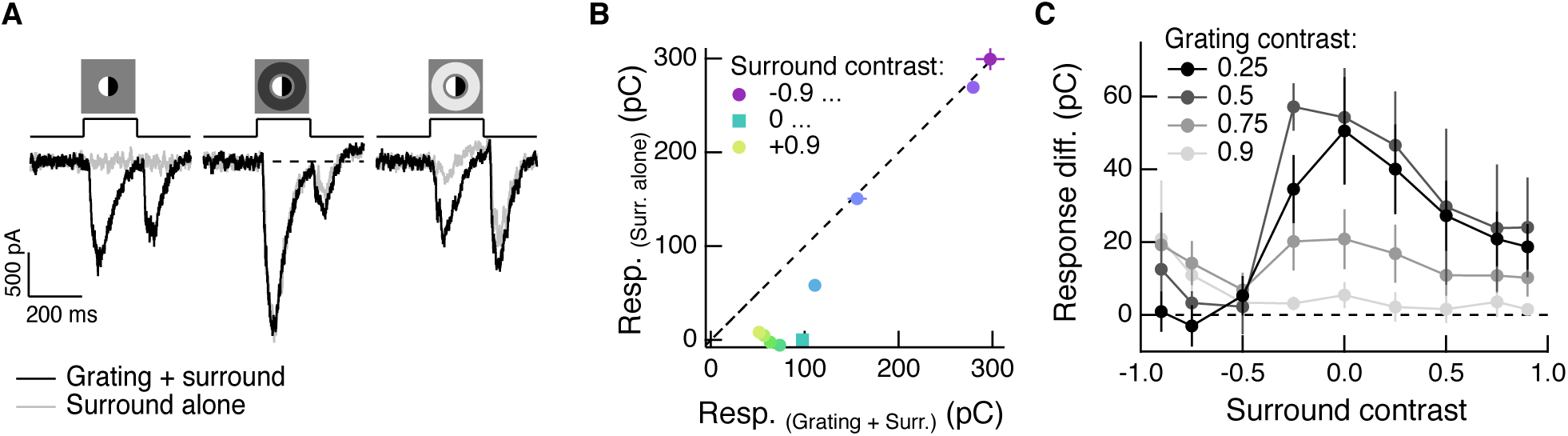
Regulation of spatial integration by the RF surround of On parasol RGCs; To accompany Fig. 3. Compare to figure 3A-C. (A) Left: On parasol RGC excitatory current response to a split-field grating stimulus in the RF center. Right: when the center stimulus is paired with a dark surround (depolarizing for this On-center RGC), the grating and the linear equivalent stimulus produce very similar excitatory current responses. (B) For the example cell in (A), we tested sensitivity to the center grating stimulus with a range of contrasts presented to the surround. Positive contrast surrounds (hyperpolarizing) decrease the response. Negative contrast surrounds (depolarizing) sum sub-linearly with the grating stimulus such that for the darkest surrounds, the addition of the grating only mildly enhances the cells response. Points show mean *±* S.E.M. excitatory charge transfer. (C) We measured the response difference between the grating stimulus and the surround-alone stimulus across a range of surround contrasts (horizontal axis) and for four different central grating contrasts (different lines). For each grating contrast, addition of either a bright or dark surround decreased sensitivity to the added grating. Points are population means *±* S.E.M. (n = 6 On parasol RGCs).

**Figure S3:**
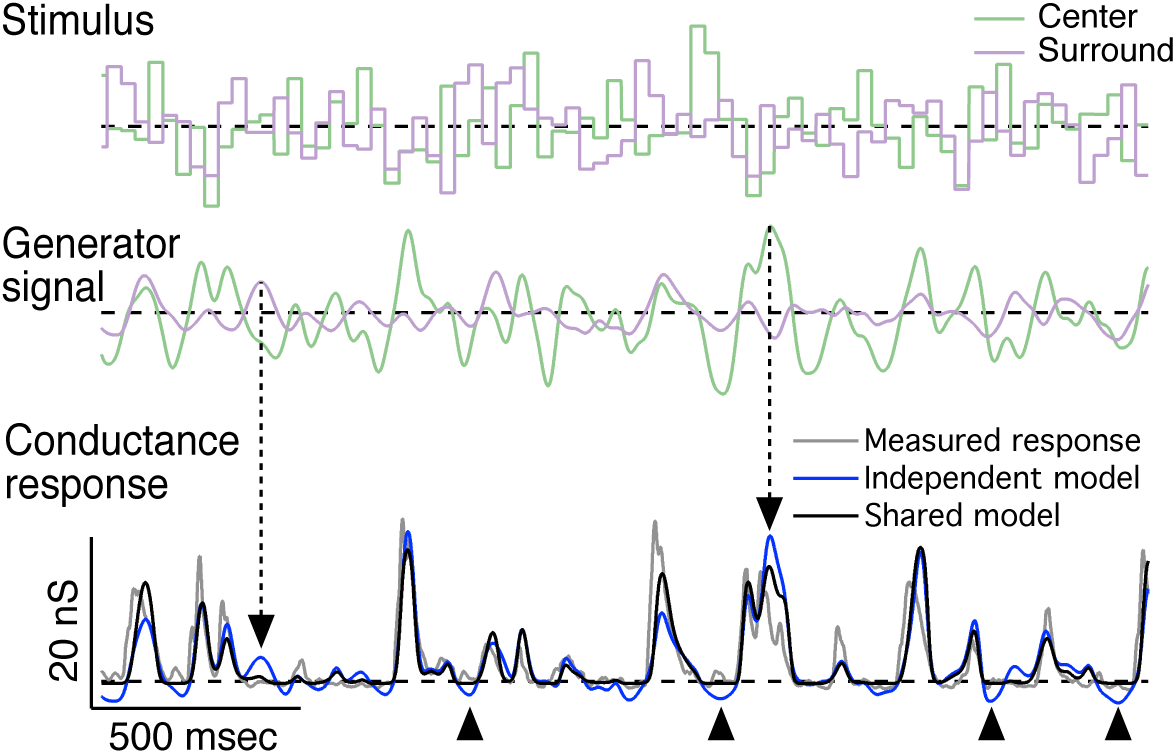
Representative predictions of linear-nonlinear cascade models for center-surround interactions; To accompany Fig. 4. Representative traces for the center-surround LN cascade models fit to the example cell in Fig. 4. Top: center-surround stimulus trace; middle: associated generator signal for the center and surround; bottom: measured conductance response (gray trace) and predicted responses from the independent nonlinearity model (blue trace) and the shared nonlinearity model (black trace). Dashed arrows indicate times at which center and surround generator signals are opposed to one another (i.e. center excited and surround inhibited or vice-versa). As a result, the independent model prediction overshoots that of the shared model and the measured response. Arrowheads below the trace indicate times at which the independent model prediction undershoots the cell’s response and the shared model prediction. These undershoots are the result of the independent model trying to account for the antagonistic relationship between the center and surround. In the independent model, the surround will suppress the response whether the center is activated or not. This is not true of the shared model or the real cell.

**Figure S4:**
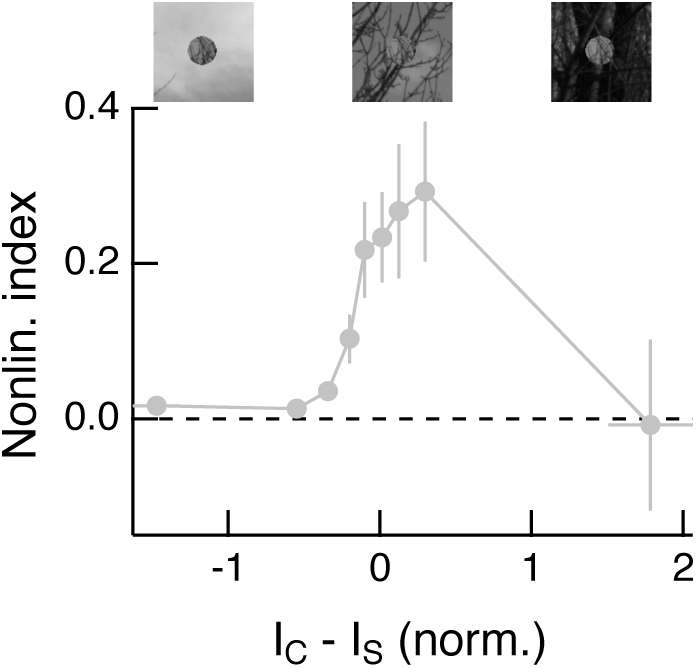
Center-surround intensity differences modulate spatial contrast sensitivity for randomly-shuffled natural surrounds; To accompany Fig. 8. From the data shown in Fig. 8D-F using the shuffled surround condition, we plotted the nonlinearity index as a function of the difference in mean intensity between the RF center and surround. Compare to Fig. 2C. Shown are mean *±* S.E.M. NLI values in equally-populated bins. Note that there are very few data points on the far right of the x axis because Off parasol RGCs very rarely spike in response to these stimuli (bright center and dark surround). Image patches above show example center & shuffled surround stimuli for the indicated center-surround intensity difference.

**Figure S5:**
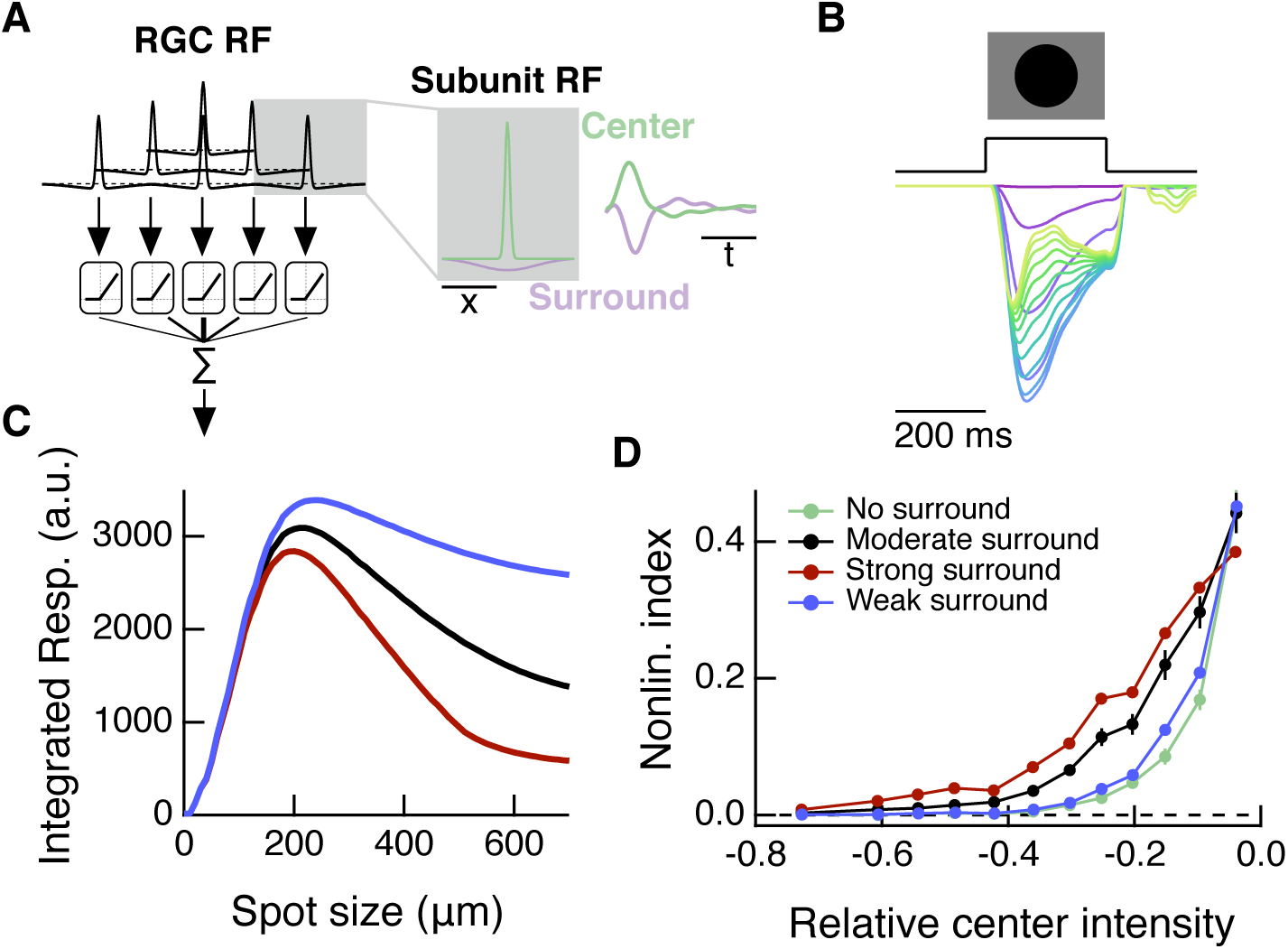
A spatiotemporal RF model also shows modulation of spatial contrast sensitivity with surround activation; To accompany Fig. 8. Compare to Fig. 8G-I, which shows results for a simpler, spatial-only RF model. (A) In addition to the spatial subunit filters, this spatiotemporal model includes temporal linear filters which were measured from excitatory current responses to white noise stimulation (Fig. 4). The temporal filters are applied to RF subunit center and surround sub-regions. (B) Response of the model (in arbitrary units) to dark spots of increasing diameter centered over the RF. From magenta to blue to yellow/green the traces show responses to spots of increasing diameter. (C) The spatiotemporal RF model shows center-surround organization to expanding spots stimuli. (D) As with the simpler spatial RF model (Fig. 8), the spatiotemporal RF model shows an increase in spatial contrast sensitivity in the face of stronger center luminance signals, and this effect depends on surround strength.

